# A Study on the Pharmacokinetic/Pharmacodynamic Profiles of the Novel PPAR Pan Agonist Chiglitazar Sodium in Rats with Hypoalbuminemia

**DOI:** 10.1101/2025.11.09.687466

**Authors:** Wenjuan He, Xinhui Zhang, Menghao Li, Bokun Chen, Yunfei Tian, Qian Sun, Xiaojuan Zhao, Yonghong Zhao, Dan Li, Xiuju Liu

## Abstract

**Objective:** To compare the changes in the pharmacokinetic/pharmacodynamic (PK/PD) profiles of Chiglitazar Sodium between healthy rats and those with hypoalbuminemia, and to evaluate the medication safety of Chiglitazar in rats with hypoalbuminemia.

**Methods:** Liquid Chromatography-Tandem Mass Spectrometry (LC-MS/MS) method was employed to determine the plasma concentration ofChiglitazar Sodium in both healthy rats and rats with hypoalbuminemia after administration. Pharmacokinetics were calculated based on the measured data. Blood glucose levels were monitored and recorded in both groups to observe pharmacodynamic changes.

**Results:** Pharmacokinetic analysis revealed that, compared to the healthy group, only the T_max_ in the hypoalbuminemia group was significantly earlier by 34.3% (P<0.001), while other pharmacokinetic parameters showed no significant differences. Pharmacodynamic studies indicated that blood glucose levels remained stable in both healthy rats and rats with hypoalbuminemia during the 0.25–10 hour monitoring period.

**Conclusion:** Except for T_max_, there were no significant differences in PK/PD parameters between rats with hypoalbuminemia and healthy rats. Blood glucose levels remained stable in both groups after oral administration of Chiglitazar Sodium. The animal experimental results suggest that, under normal conditions, hypoalbuminemia does not affect the pharmacokinetic and pharmacodynamic profiles of Chiglitazar Sodium.

---

Chiglitazar Sodium is a novel PPAR pan-agonist with independent intellectual property rights in China^[1]^. It works by activating PPARα, PPARγ, and PPARδ receptors to enhance insulin sensitivity, regulate blood glucose levels, and promote fatty acid oxidation and utilization. It is clinically used for the treatment of type 2 diabetes and metabolic regulation^[2]^. Hypoalbuminemia is a common complication in patients with type 2 diabetes, particularly among the elderly and those with poor glycemic control^[3]^. Under normal conditions, drugs bind to plasma proteins at a certain ratio, with the free drug fraction being pharmacologically active^[4]^. However, patients with hypoalbuminemia have reduced plasma protein content, which can directly affect pharmacokinetic processes^[5]^. Chiglitazar Sodium is a highly protein-bound drug^[6]^. Due to its relatively recent market approval, the drug’s prescribing information lacks safety data for use in special populations. This study was designed to evaluate the safety of Chiglitazar in rats with hypoalbuminemia by comparing changes in PK/PD parameters between healthy rats and those with hypoalbuminemia, thereby laying the groundwork for further clinical research on the use of Chiglitazar Sodium in patients with hypoalbuminemia.

## 1 Materials and Methods

### 1.1 Durgs and chemicals

The reference standard ofChiglitazar Sodium (Batch No. 101548-202101) was obtained from the National Institutes for Food and Drug Control. Chiglitazar Sodium Impurity-A (Batch No. 20211026) and Chiglitazar Sodium Tablets (Batch No. 221001) were supplied by Shenzhen Chipscreen Biosciences Co., Ltd. Doxorubicin Hydrochloride (Batch No. 20000866-DX-315-A22071102) was provided by Shijiazhuang Pharmaceutical Group Ou Yi Pharmaceutical Co., Ltd. Heparin Sodium Injection (Batch No. 230803-1) was sourced from Ma’anshan Fengyuan Pharmaceutical Co., Ltd. Chromatographic grade methanol and acetonitrile (Batch Nos. F24O43201 and F24O7A209) were purchased from Thermo Fisher Scientific.

### 1.2 Instruments

The experimental work was performed using the following equipment: a liquid chromatography-mass spectrometry system (LCMS-8040CL, Shimadzu, Japan), a C8 chromatographic column (5262-0032, Phenomenex), a centrifuge (ABBOTT, Germany), a vortex mixer (XW-80A, Shanghai Medical University Instrument Factory), a low-temperature freezer (MDF-U2086S, Sanyo, Japan), a refrigerator (YC-180, Aucma), and an electronic analytical balance (CPA225D, Sartorius, Germany).

### 1.3 Analytical Conditions

#### 1.3.1 Chromatographic Conditions

A Phenomenex C8 column protected by a C8 guard column was utilized under the following conditions: a mobile phase of water (A) and acetonitrile (B) (17:83, v/v) in an isocratic elution mode, a flow rate of 0.35 mL/min, a column temperature of 40 °C, and an injection volume of 5 μL.

#### 1.3.2 Mass Spectrometric Conditions

The mass spectrometric detection was conducted on an LC-MS/MS system under the control of Labsolution software. The system was operated in negative ionization mode. The monitoring ions were set as m/z 571.2→527.2 for chiglitazar and m/z 553.2→509.3 for IS.

### 1.4 Animals

Specific pathogen-free (SPF) male Sprague-Dawley (SD) rats, aged 5 weeks with an initial body weight of 170–180 g, were obtained from Beijing Huafukang Bioscience Co., Ltd. (Laboratory Animal License No.: SCXK (Jing) 2024-0003). The study protocol was approved by the Institutional Animal Care and Use Committee, and all procedures were conducted in accordance with the approved guidelines (Ethical Approval No.: 2024-AE336). The rats were housed under standard laboratory conditions at room temperature with a relative humidity of 40%–60% and were acclimatized for one week prior to experimentation.

### 1.5 Solution Preparation

#### 1.5.1 Preparation of Standard Solutions

Chiglitazar Sodium and the internal standard reference substances were accurately weighed (1 mg each), dissolved in methanol, and diluted to volume to prepare stock solutions with a concentration of 100 μg/mL. The solutions were stored at −40°C for future use.

#### 1.5.2 Preparation of the Gavage Solution

The administered dose for rats was 3.33 mg/kg (calculated based on the dosage conversion from the Chiglitazar Sodium prescribing information [7]). One tablet of Chiglitazar Sodium was placed in a beaker, and 16 ml of purified water was added. The mixture was stirred until the tablet disintegrated and dissolved completely, resulting in a gavage solution with a concentration of 10 mg/ml. The solution was freshly prepared before each administration.

#### 1.5.3 Preparation of the Modeling Agent Injection

Doxorubicin hydrochloride (20 mg) was accurately weighed and dissolved in 0.9% sodium chloride injection to prepare a doxorubicin hydrochloride injection with a concentration of 2 mg/ml. The injection was freshly prepared before modeling.

### 1.6 Methodological Validation

The analytical method in this study was established with reference to the LC-MS/MS method for determining Chiglitazar Sodium described by N. N. Chu et al. [9]. The signal-to-noise ratio (S/N) of the instrument was 11.2 at 8 ng/mL, and the lower limit of quantitation was set at 10 ng/mL. The quantitative range of Chiglitazar Sodium was defined as 10–2000 ng/mL for method validation.

Calibration standards were prepared at concentrations of 10, 50, 100, 500, 1000, and 2000 ng/mL in simulated plasma to construct a standard curve. Quality control (QC) samples at low, medium, and high concentrations (10, 500, and 2000 ng/mL) were processed and analyzed according to the procedure described in Section 1.8. Each concentration was prepared in five replicates. The relative recovery and intra-day precision were calculated based on the ratio of the measured concentration to the theoretical concentration. Inter-day precision was assessed by analyzing five replicates at each concentration over three consecutive days. Stability was evaluated under the following conditions: QC samples were kept at room temperature for 8 h, stored at –40°C for 7 days, and subjected to three freeze–thaw cycles at –40°C. After treatment, samples were processed and analyzed as described in Section 2.3 to investigate placement stability.

### 1.7 Modeling of Hypoproteinemia in Rats

After a 7-day adaptation period, 16 male Sprague-Dawley (SD) rats were divided into two groups, with 8 rats in each group. Rats in the experimental group were administered doxorubicin hydrochloride at a dose of 7.5 mg/kg via a single tail vein injection[7]. Rats in the control group were administered a corresponding volume of 0.9% sodium chloride injection based on body weight, also via a single tail vein injection. Fifteen days after the injection, 1.5 ml of blood was collected from the inner canthus of both eyes ofthe rats in each group using sterile serum separator gel coagulant tubes. The serum total protein and albumin levels of the rats were measured at the Clinical Laboratory of the Second Hospital of Hebei Medical University. A serum total protein level below 50 g/L and a serum albumin level below 25 g/L indicated successful modeling of hypoproteinemia in rats [8]. The baseline data of the rats involved in the experiment are shown in Table 1.

**Table 1.** Baseline characteristics of rats are presented as mean ± SD (n=8)

| Characteristics (mean±SD) | | Healthy rats |
| --- | --- | --- |
| Hypoalbuminemic rats |  |  |
| body weight (g) | 182.2±3.10 | 203.6±5.30 |
| total protein (g/l) | 43.61±3.56 | 61.07±3.71 |
| albumin (g/l) | 19.7±2.75 | 31.6±1.30 |

### 1.8 Plasma Sample Processing

A precise volume of 10 μL ofthe internal standard working solution (5000 ng/mL) was pipetted into a 1.5 mL centrifuge tube and evaporated to dryness under a stream of nitrogen. Then, 100 μL of plasma sample was added and vortex-mixed for 1 minute. Subsequently, a threefold volume of acetonitrile:methanol:water (90:7:3, v/v/v) solution was added to the plasma sample to precipitate proteins. After vortex mixing for 1 minute, the mixture was centrifuged at 10,900 r/min for 10 minutes. The resulting supernatant was collected for injection and analysis.

### 1.9 Drug Administration and Blood Sample Collection

After fasting for 10 hours, SD rats in both groups were orally administered Chiglitazar Sodium at a dose of 3.33 mg/kg. Blood samples (0.5 mL) were collected from the inner canthus at 0, 0.25, 0.5, 1, 1.5, 2, 3, 4, 5, 6, 8, 10, 12, and 24 hours post-dosing and placed in heparinized centrifuge tubes. Additionally, blood samples (0.3 mL) were collected at 0, 0.25, 0.5, 1, 1.5, 2, 3, 4, 5, 6, 8, and 10 hours for blood glucose measurement using a glucometer. Blood samples at verlapping time points were collected only once. After the 10-hour blood glucose collection, the rats were allowed free access to water and food. Following collection, PK blood samples were centrifuged at 10,900 r/min for 5 minutes to separate the plasma, which was then stored at −40°C pending further analysis.

## 2 Results

### 2.1 Specificity

The retention times for both Chiglitazar Sodium and the internal standard (Chiglitazar Sodium Impurity-A) were 0.7 min. The results demonstrate that the developed method exhibits high specificity, with no interference from endogenous substances in rat plasma on the analyte quantification. This method is suitable for the determination of blood concentration ofChiglitazarSodium. A representative chromatogram is shown in Figure 1.

**Figure 1.**
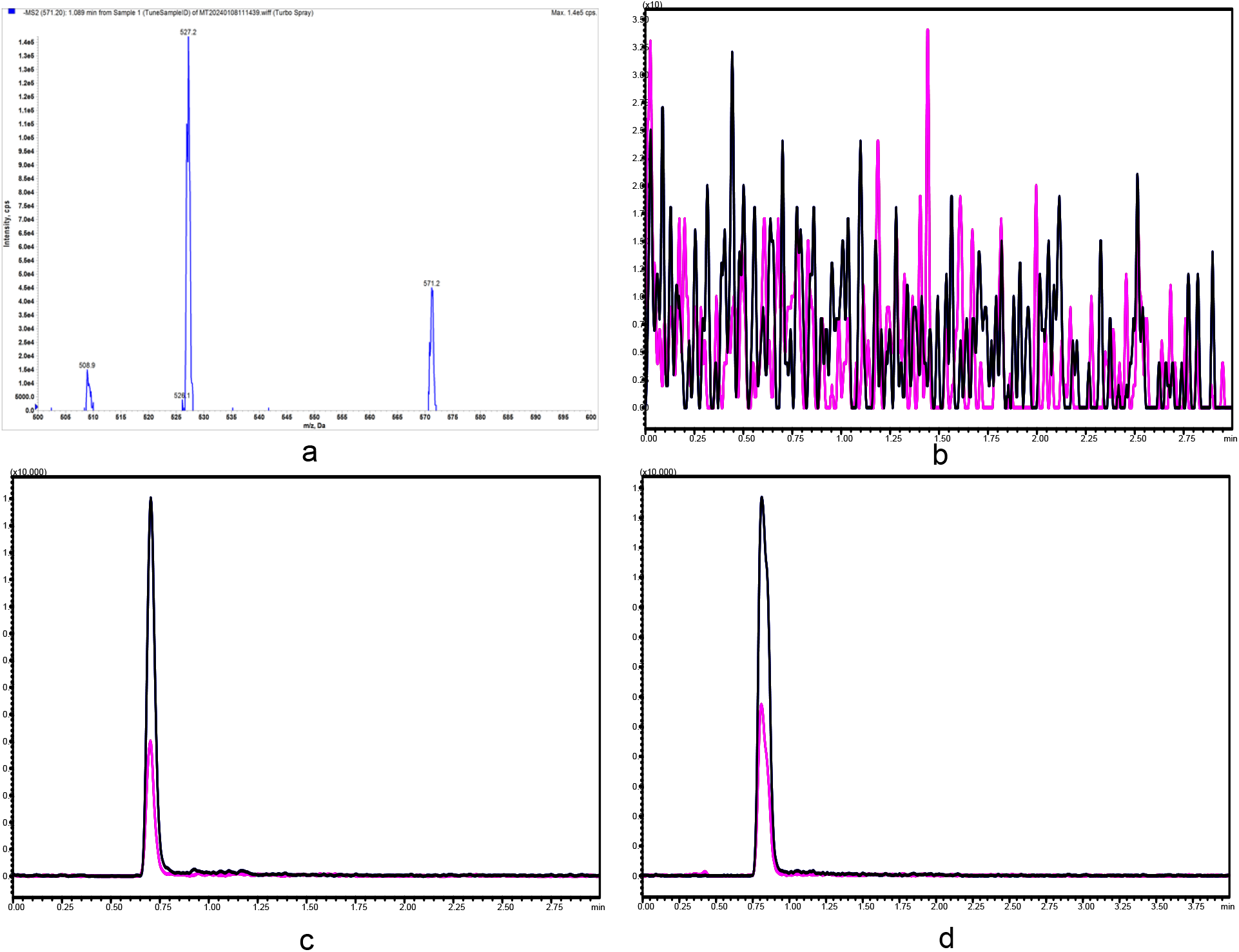
LC-MS/MS Chromatogram ofChiglitazar Sodium. Note: (a) Precursor-product ion spectrum of Chiglitazar Sodium; (b) Chromatogram of blank plasma; (c) Chromatogram of spiked plasma sample; (d) Chromatogram of rat plasma sample. The black curve represents Chiglitazar Sodium; th e red curve represents the internal standard.

### 2.2 Standard Curve

The linear range for Chiglitazar Sodium was 10 ng/mL to 2000 ng/mL. The standard curve equation was Y = 0.834381X+0.0113496 (r^2^ = 0.9998), demonstrating excellent linearity. The lower limit of quantification (LLOQ) was 10 ng/mL.

### 2.3 Precision and Accuracy

The intra-day and inter-day relative standard deviations (RSD) for Chiglitazar Sodium did not exceed 10%. The results indicate that the method meets the acceptance criteria for bioanalytical method validation and demonstrates good reproducibility. Detailed data are summarized in Table 2.

**Table 2.** Intra-day and Inter-day Precision of Chiglitazar Sodium in Rat Plasma(*n*=5)

| Concentration<br>(ng/mL) | Intra-day Precision |  |  | Inter-day Precision |  |  |
| --- | --- | --- | --- | --- | --- | --- |
|  | Mean±SD | RSD | RE | Mean±SD | RSD | RE |
|  |  | (%) | (%) |  | (%) | (%) |
| 10 | 10.21±0.59 | 5.80 | 105.6 | 10.22±0.618 | 6.05 | 105.4 |
| 500 | 509.96±4.98 | 0.98 | 102.0 | 503.4±7.842 | 1.56 | 101.4 |
| 1000 | 1977.8±73.37 | 3.71 | 103.4 | 2050±39.98 | 1.95 | 102.8 |

### 2.4 Recovery and Matrix Effect

The extraction recovery ofChiglitazar Sodium exceeded 90%, with an RSD of less than 10%. The matrix effect fell within ±10%, RSD < 10%. These results demonstrate that the method complies with accepted criteria for bioanalytical method validation (e.g., FDA/EMA guidelines) and exhibits good reproducibility. Detailed data are presented in Table 3.

**Table 3.** Recovery and Matrix Effect of Chiglitazar Sodium in Rat Plasma(*n*=5)

| Concentration<br>(ng/mL) | Recovery |  | Matrix Effect |  |
| --- | --- | --- | --- | --- |
|  | Mean±SD | RSD (%) | Mean±SD | RSD (%) |
| 10 | 103.98±3.37 | 3.24 | 102.14±6.50 | 6.36 |
| 500 | 101.91±2.58 | 2.53 | 100.57±2.75 | 2.73 |
| 1000 | 102.03±2.46 | 2.41 | 101.5±4.02 | 3.96 |

The results indicated that the RSD of plasma samples remained below 15% after being stored at room temperature for 8 hours, at −40 °C for 7 days, and following three freeze-thaw cycles.

These findings demonstrate that Chiglitazar Sodium exhibits good stability under the aforementioned conditions, as summarized in Table 4.

**Table 4.**
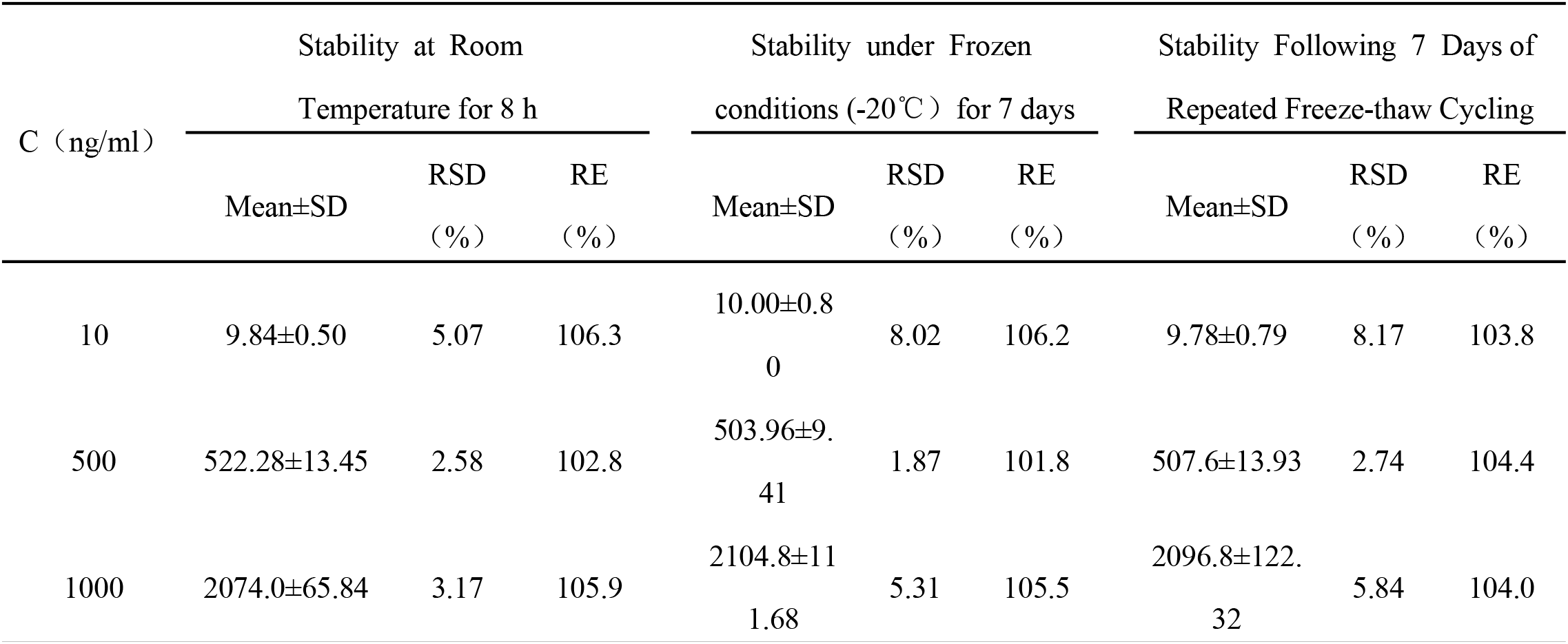
Stability of Chiglitazar Sodium in Rat Plasma(*n*=5)

| C (ng/ml) | Stability at Room |  |  | Stability under Frozen |  |  | Stability Following 7 Days of |  |  |
| --- | --- | --- | --- | --- | --- | --- | --- | --- | --- |
|  | Temperature for 8 h |  |  | conditions (-20℃ ) for 7 days |  |  | Repeated Freeze-thaw Cycling |  |  |
|  | Mean±SD | RSD (%) | RE (%) | Mean±SD | RSD (%) | RE (%) | Mean±SD | RSD (%) | RE (%) |
| 10 | 9.84±0.50 | 5.07 | 106.3 | 10.00±0.80 | 8.02 | 106.2 | 9.78±0.79 | 8.17 | 103.8 |
| 500 | 522.28±13.45 | 2.58 | 102.8 | 503.96±9.41 | 1.87 | 101.8 | 507.6±13.93 | 2.74 | 104.4 |
| 1000 | 2074.0±65.84 | 3.17 | 105.9 | 2104.8±111.68 | 5.31 | 105.5 | 2096.8±122.32 | 5.84 | 104.0 |

The mean plasma concentration-time curve of Chiglitazar Sodium in rats following administration (3.33 mg/kg) is shown in Figure 2. The pharmacokinetic parameters of rats in the control and experimental groups after administration are summarized in Table 5.

**Table 5.** Pharmacokinetic Parameters ofChiglitazar Sodium in Hypoalbuminemic and Healthy Rats after Administration(n=8)

| Parameters | Hypoalbuminemic | Healthy rats | P value |
| --- | --- | --- | --- |
| AUC<sub>0-t</sub> (ng/L/h) | 3092.125±439.934 | 2819.407±588.956 | 0.312 |
| AUC<sub>0-∞</sub> (ng/L/h) | 3485.747±589.851 | 3000.148±584.800 | 0.120 |
| t<sub>1/2</sub> (h) | 6.994±2.283 | 5.164±2.215 | 0.126 |
| T<sub>max</sub> (h) | 2.875±0.641 | 4.5±0.535 | <0.001 |
| V<sub>d/F</sub> (L/kg) | 9.64±2.707 | 8.594±3.947 | 0.547 |
| CL/F (L/h/kg) | 1.835±1.427 | 2.233±1.009 | 0.530 |
| C<sub>max</sub> (ng/L) | 403.677±100.917 | 454.944±71.822 | 0.085 |

**Figure 2.**
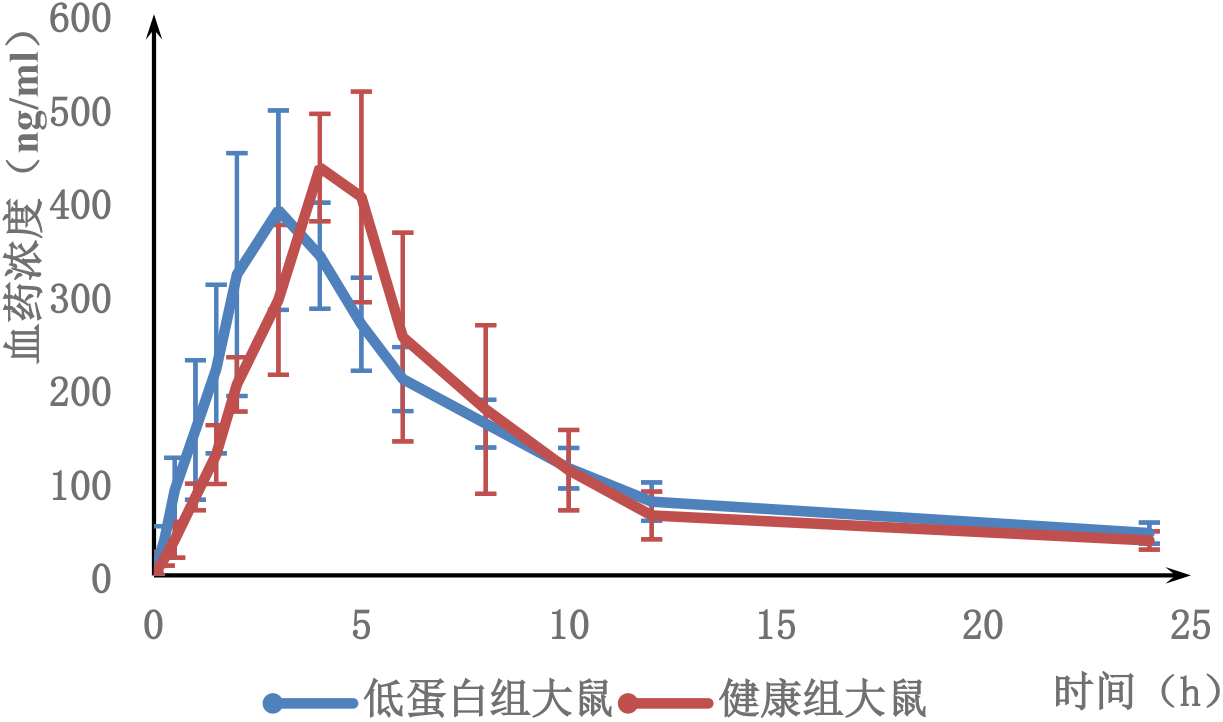
Mean Plasma Concentration-Time Profile of Chiglitazar Sodium in Hypoalbuminemic versus Healthy Rats following a Single Oral Dose(n=8)

Pharmacokinetic analysis revealed that compared with the healthy group, the hypoproteinemia group showed a significantly earlier Tmax, with a reduction of 36.11% (P < 0.001). In hypoproteinemic rats, AUC0–t (mg/L·h), AUC0–∞ (mg/L·h), t1/2 (h), and Vd/F increased by 9.67%, 16.18%, 35.43%, and 12.17%, respectively, compared with healthy rats, these differences were not statistically significant (P > 0.05). Meanwhile, Cmax and CL/F decreased by 11.26% and 17.82% in hypoproteinemic rats compared with healthy rats, also without significant differences (P > 0.05).

Blood glucose data collected from 0 to 10 hours are presented in Figure 3. The value at 0 h represents the fasting blood glucose level after 10 hours of fasting, while data from 0.25 to 10 h correspond to blood glucose levels after administration of Chiglitazar Sodium in both hypoproteinemic and healthy groups. The average fasting blood glucose was 4.46 mmol/L in healthy rats and 4.84 mmol/L in hypoproteinemic rats. After drug administration, blood glucose levels remained stable from 0.25 to 10 h in both groups, with mean values of 6.63 mmol/L in the healthy group and 5.74 mmol/L in the hypoproteinemia group. The difference in average blood glucose between the two groups was not statistically significant (P = 0.084).

**Figure 3.**
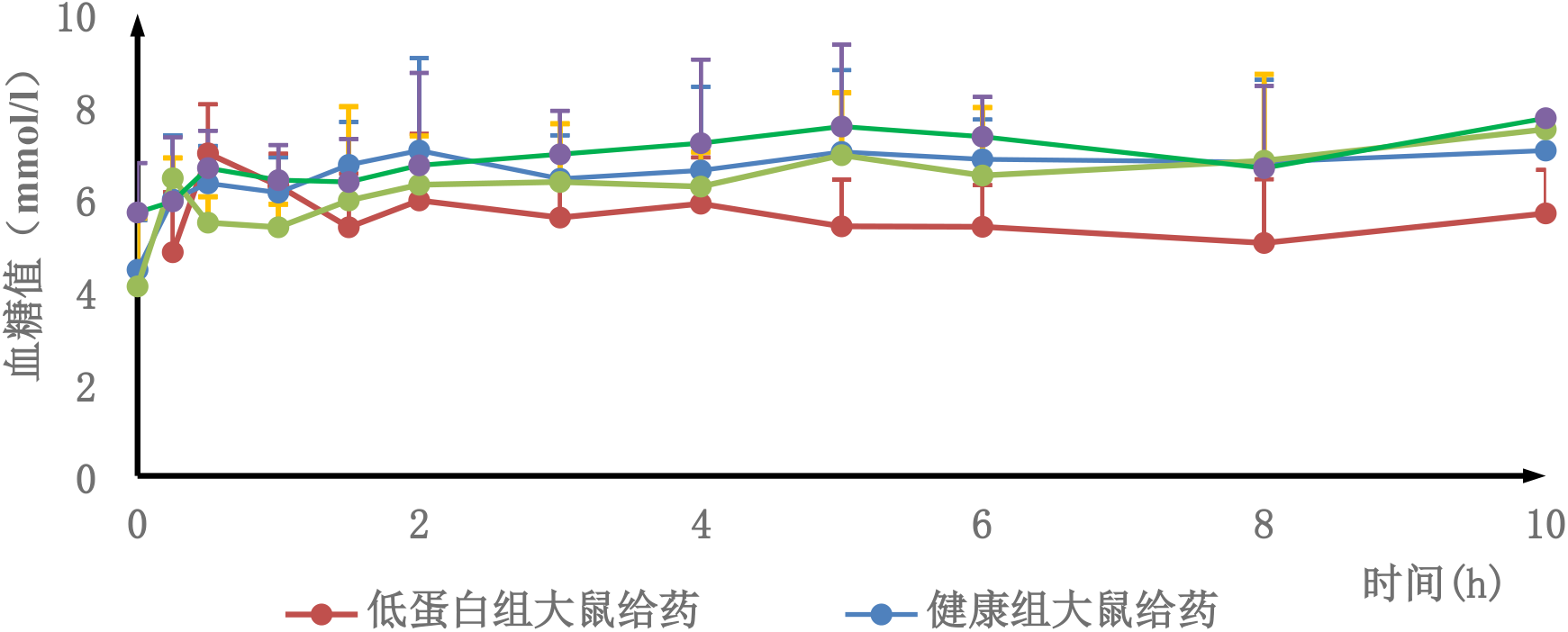
Blood Glucose Levels in Hypoalbuminemic and Healthy Rats with and without Chiglitazar Sodium Administration(n=8)

## 3 Discussion

This study revealed certain differences in the PK/PD characteristics of Chiglitazar Sodium at a dose of 3.33 mg/kg between hypoproteinemic and healthy rats. Pharmacokinetic data showed that the Tmax in the hypoproteinemia group occurred 36.11% earlier than that in the healthy group (P < 0.001), indicating a statistically significant difference. Tmax primarily reflects the rate of drug absorption into the systemic circulation; a faster absorption rate results in a shorter time to peak concentration [6–9]. Chiglitazar Sodium is absorbed into the bloodstream via passive transport through the gastrointestinal tract [10–11]. After entry into the circulation, the free drug binds to plasma proteins, forming a protein-bound fraction. This binding reduces the concentration of free drug in the blood, thereby maintaining a concentration gradient of the free drug between the intestinal lumen and the blood, which further facilitates passive diffusion of the drug from the gastrointestinal tract into the circulation [12–15]. Hypoproteinemia may reduce drug-protein binding, increase the free drug concentration, and consequently accelerate distribution and metabolism, leading to a shorter Tmax [13–15]. Furthermore, decreased plasma colloidal osmotic pressure and reduced plasma volume in hypoproteinemic rats may promote drug distribution to some extent, which could explain the somewhat lower Cmax observed in the hypoproteinemia group compared with the healthy group, although the difference was not statistically significant (P > 0.05). The doxorubicin used in the modeling process may have caused certain degrees of hepatic and renal impairment in the rats. Combined with reduced colloidal osmotic pressure and decreased renal blood flow, this injury may affect drug metabolism and excretion, manifesting as increases in AUC0–t, AUC0–∞, and t1/2 in the hypoproteinemia group compared with the healthy group (P > 0.05). However, aside from Tmax, none of the other pharmacokinetic parameters showed significant differences between the two groups. This may suggest that hypoproteinemia has limited impact on the distribution, metabolism, and excretion of Chiglitazar Sodium, although further investigation is required to elucidate the underlying mechanisms [12–15].

Pharmacodynamic data indicated that the blood glucose profiles of both rat groups remained relatively stable and did not exhibit significant fluctuations corresponding to the changes in plasma drug concentration. Furthermore, there was no statistically significant difference in blood glucose levels between the hypoproteinemia group and the healthy group (P > 0.05). According to literature reports, the normal blood glucose range in rats is 4.0–6.5 mmol/L [16]. As shown in Figure 3, most blood glucose measurements in each group fluctuated within this normal range, suggesting that Chiglitazar Sodium did not produce a significant glucose-lowering effect in healthy rats [17–18]. The observation that some rats exhibited elevated blood glucose levels above normal fasting values after administration may be attributed to stress responses induced by experimental procedures such as oral gavage and blood collection from the inner canthus [19–20]. Such stress responses can activate the sympathetic nervous system and the hypothalamic-pituitary-adrenal (HPA) axis, leading to the release of stress hormones and consequent hyperglycemia [21– 22].

During the experiment, one hypoglycemic event was observed in each of the healthy group and the hypoproteinemia group. In the hypoproteinemia group, Rat #7 exhibited blood glucose levels of 3.9 mmol/L and 3.2 mmol/L at 6 and 8 hours after administration, respectively. However, without intervention, its blood glucose returned to 5.3 mmol/L after 10 hours. Furthermore, Rat #7 in the hypoproteinemia group showed a 32.6% higher Cmax (P = 0.052) and a 24.01% higher AUC0–∞ (P = 0.29) compared to the average values of the other seven rats in the same group, indicating enhanced drug absorption relative to its counterparts. In the healthy group, Rat #6 also experienced a hypoglycemic event at 10 hours post-dosing, with a blood glucose level of 3.8 mmol/L. This rat demonstrated a 26.4% higher C_max_ (P = 0.092) and a 48% higher AUC_0–t_ (P=0.071) compared to the average of the other seven rats in the group, similarly suggesting greater drug absorption capacity. Since all hypoglycemic events occurred after 16 hours of fasting, it is hypothesized that the occurrence of hypoglycemia may be associated with individual variability and prolonged fasting.

Also, this study has several limitations that should be acknowledged: 1. Experimental stress in rats: We conducted validation experiments and observed that oral gavage administration—even using purified water—elevated blood glucose levels in both healthy and hypoproteinemic groups. Previous literature also indicates that stress during experimental procedures can increase blood glucose levels, even under glucose-lowering treatment, potentially masking drug-induced effects [20]. Therefore, the observed increases in blood glucose in the pharmacodynamic evaluations are likely stress-related and not correlated with the pharmacological activity of Chiglitazar Sodium. 2. Undetected hepatic and renal function: The animal model involved male SD rats treated with 7.5 mg/kg doxorubicin to induce hypoproteinemia. This agent is known to cause hepatorenal toxicity, which may alter drug metabolism and excretion. However, hepatic and renal functions were not explicitly monitored, limiting comprehensive interpretation of the pharmacokinetic results. 3. Species and disease-state differences: Translating findings from rats to humans is constrained by interspecies variations. Furthermore, real-world patients exhibit diverse disease states and comorbidities, which may significantly influence drug metabolism and response in ways not captured in this experimental model.

## 4 Conclusion

In summary, the hypoproteinemia group demonstrated a significant difference only in Tmax compared with healthy rats. Under typical circumstances, Chiglitazar Sodium can be safely administered in patients with hypoproteinemia, though individual variability should be considered.

Since Chiglitazar Sodium is primarily metabolized by the liver, and the prescribing information indicates that moderate hepatic impairment increases Cmax and AUC by 17% and 74%, respectively, further validation of its safety is warranted in hypoproteinemic patients with concomitant liver dysfunction.

## 5 Limitations

This study utilized male Sprague-Dawley rats in which hypoproteinemia was induced using doxorubicin at a dose of 7.5 mg/kg. It should be acknowledged that doxorubicin may cause hepatorenal toxicity, which could influence drug metabolism and pharmacokinetics. However, no direct assessment of hepatic or renal function was performed following successful model establishment.

Furthermore, blood glucose levels can be influenced by multiple factors. The stress response induced by oral gavage—coupled with individual variability in stress reactivity among rats—may have confounded glycaemic measurements. In contrast, human patients typically exhibit better compliance with oral medications and are less prone to procedure-induced hyperglycemia.

Additionally, interspecies differences between rodents and humans, along with variations in clinical disease states among patients, may lead to divergent drug metabolism profiles. Therefore, further validation is warranted to comprehensively evaluate the safety of Chiglitazar Sodium in hypoproteinemic patients, particularly those with concurrent hepatic or renal impairment.

## Data Availability

All data supporting the findings of this study are included in the Supplementary Information of this article. Further reasonable requests can be directed to the corresponding author.

## Acknowledge

I confirm that anyone listed under the Acknowledgements section of the manuscript has been informed of their inclusion and approve this. This work was supported by Medical Science Research Project of Hebei (20241323).

## Conflict of Interests

The authors declare that there are no conflict of interests.

## Animal Ethics

I confirm the study has received approval from the ethics committee at the institution. (2024-AE336)

## Notes

### Competing Interest Statement

The authors have declared no competing interest.

